# Dynamic response to initial stage blindness in visual system development

**DOI:** 10.1101/116590

**Authors:** Erping Long, Xiayin Zhang, Zhenzhen Liu, Xiaohang Wu, Xuhua Tan, Duoru Lin, Qianzhong Cao, Jingjing Chen, Zhuoling Lin, Dongni Wang, Xiaoyan Li, Jing Li, Jinghui Wang, Wangting Li, Haotian Lin, Weirong Chen, Yizhi Liu

**Affiliations:** State Key Laboratory of Ophthalmology, Zhongshan Ophthalmic Center, Sun Yat-sen University, Guangzhou 510060, China

**Keywords:** Visual system development, plasticity, sensitive periods, retina malleability

## Abstract

Sensitive periods and experience-dependent plasticity have become core issues in visual system development. Converging evidence indicates that visual experience is an indispensable factor in establishing mature visual system circuitry during sensitive periods and the visual system exhibits substantial plasticity when facing deprivation. The mechanisms that underlie the environmental regulation of visual system development and plasticity are of great interest but need further exploration. Here, we investigated a unique sample of human infants who experienced initial stage blindness (beginning at birth and lasting 2 to 8 months) before the removal of bilateral cataracts. Retinal thickness, axial length, refractive status, visual grating acuity and genetic integrity were recorded during the preoperative period or at surgery, and then during follow-up. The results showed that the development of the retina is malleable and associated with external environment influences. Our work supported that the retina might play critical roles in the development of the experience-dependent visual system and its malleability might partly contribute to the sensitive period plasticity.

**SUMMARY STATEMENT:** The follow-up investigation of a group of human infants, who experienced initial stage blindness before the removal of bilateral cataracts, revealed that retinal development is associated with environment influences and its malleability might be a potential basis of plasticity.

## INTRODUCTION

Visual experience from external environment is crucial to the development of the entire visual system (Lewis and Maurer, 2005; Hooks and Chen, 2007). Previous evidence supported that abnormal visual experience causes dramatic functional deficits, but visual system can retain its plasticity and has potential to recover, at least in part, after visual deprivation during or even beyond classical sensitive period in adulthood (Maurer et al., 1999; Morishita and Hensch, 2008; Jeon et.al., 2012). Therefore, the mechanisms that underlie the regulation of visual system sensitive periods are currently of great interest. Although evidence from researches revealed that sensitive periods of the visual cortex are activated by distinct mechanisms (Hooks and Chen, 2007), insufficient attention has been paid to other main components of the visual system before and following initial stage blindness (Mutti et al., 2009; Wang et al., 2015), such as eyeball development. The underlying mechanisms of both the environmental regulation of visual system development and plasticity need further understandings.

Up to now, three main factors have been thought to hinder research progress on the environmental regulation of visual system development and its plasticity. First, although evidence obtained from natural deprivation models occurring in human has contributed direct understanding of cortex plasticity (Heering et al., 2016; Grady et al, 2014; Guerreiro et al., 2016), human-level investigations of environmental effects on retina and other components of visual system are still limited. The results of animal studies cannot necessarily be generalized to human and may even differ from the results obtained from human (Smith et al., 2014). Second, the evaluation metrics of development often fail to assess the entire visual system. In addition to the development of visual cortex, different parts of the eyeball often present distinct developmental patterns (Wallman and Winawer, 2004; Liu et al., 2007; Qin et al., 2013; Schaeffel and Wildsoet, 2013); therefore, it is important to integrate them to explore the experience-dependent plasticity of the visual system as a whole. Third, genetic effects are difficult to be excluded from the analysis. Previous studies have shown that genes play a key role in the visual development, as any deficiency of these genes may lead to varying degrees of visual dysplasia (Azuma et al., 1999; Barbieri et al., 1999; Hallonet et al., 1999; Minoshima et al., 1999; Lundwall et al., 2015), presenting a challenge for determining the genetic influence on visual development.

In this study, we followed a group of human infants who experienced initial stage blindness (beginning at birth and lasting 2 to 8 months) before the removal of bilateral cataracts. Having access to this rare population provides a unique opportunity to investigate the effect of form-deprivation on human visual system development. In addition, we examined four critical indicators, namely, retinal thickness (RT), axis length (AL), refractive status and visual grating acuity, to systematically investigate visual system development and its plasticity during the sensitive period. Here, the development of RT is accompanied by fluctuations in retinal functions: receiving and translating visual information in the visual pathway. We can infer the development of retinal function via RT measurement, due to the significant correlation between retinal structure and retinal cell function (Rolle et al., 2016). Measurement of AL and refractive status can provide valuable assays for the dynamic emmetropization process after birth (Pennie et al., 2001). Moreover, grating acuity may be recorded as a valuable metric for the overall visual assessment of infants and young children (Skoczenski and Norcia, 1999). All included children had no family history of visual impairment to mimic the environmental manipulation processes following initial stage blindness. Whole exome sequencing was used to identify all of the potential genomic deficiencies associated with visual development. The present work may serve as a valuable reference for future studies of visual system development and provides a fresh paradigm for understanding the developmental process from the clinicians’ perspective.

## RESULTS

### Longitudinal assessment protocol

The study pipeline is presented in Figure 1. Four critical indicators were measured: 1) RT for functional development of the retina for receiving and translating visual information, 2) AL for investigating holistic eye emmetropization during initial stage blindness, 3) refractive status for the dynamic emmetropization process after surgery, and 4) visual grating acuity for the assessment of the overall visual system development. In addition, we use whole exome sequencing to determine whether the children had genomic deficiencies for visual development.

**Figure 1.**
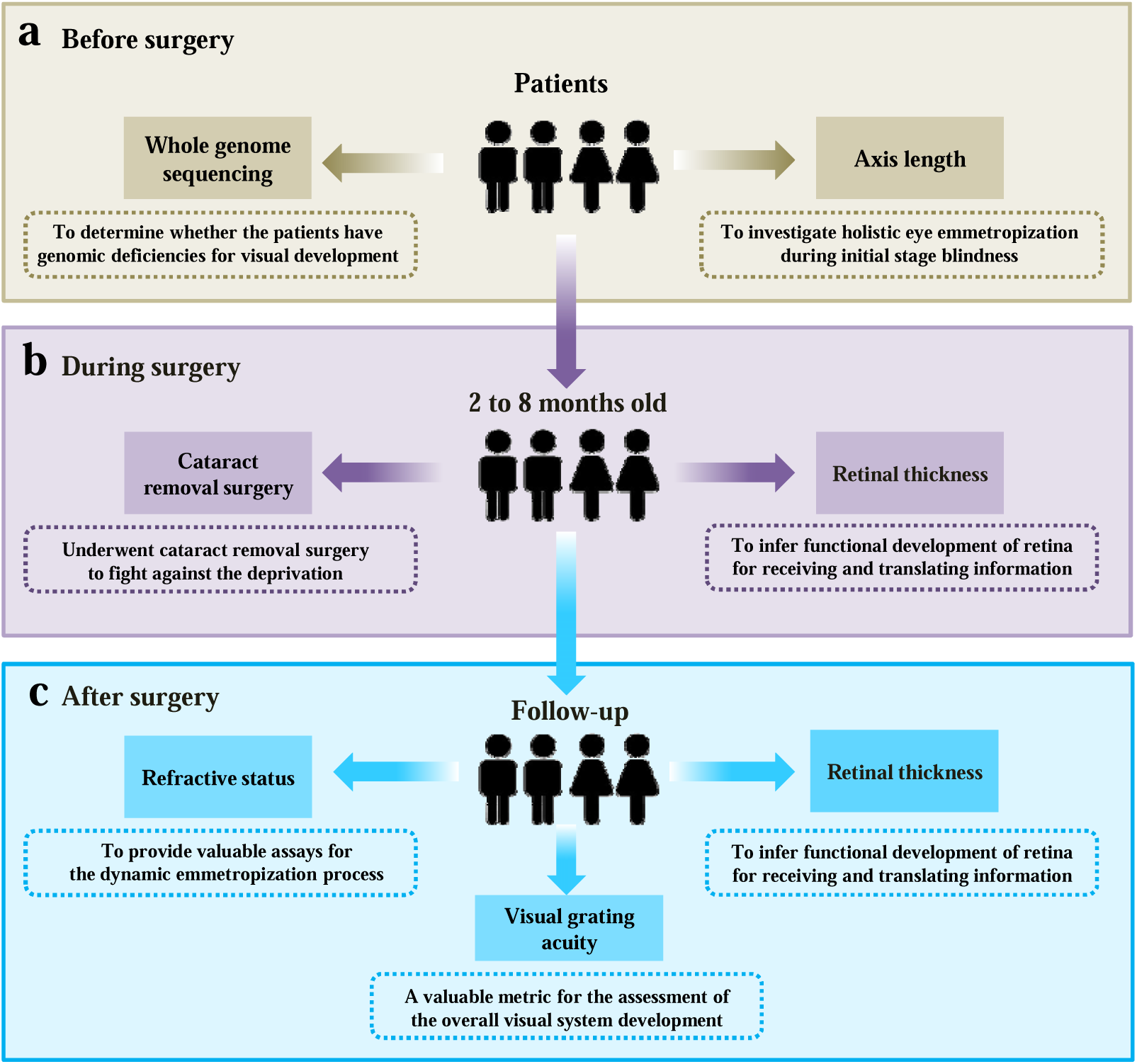
Pipeline for the investigation of initial stage blindness in children. **a,** AL measurements and whole exome sequencing were conducted prior to surgery. **b,** The first RT measurement was conducted immediately following cataract removal during the surgery. **c,** Longitudinal assessments consisting of RT, refraction and VA measurements were conducted postoperatively at 1 week, 1 month, 3 months, 6 months and every 6 months thereafter. (Notes: AL=axial length; RT=retinal thickness; VA=visual acuity).

AL measurements and baseline visual acuity (VA) evaluations were conducted before surgery (Figure 1a) (Pennie et al., 2001; Mutti et al., 2005). The dense and total cataracts hindered their preoperative RT measurement; consequently, the first RT measurement was conducted immediately following the cataract removal during the surgery (Figure 1b). Longitudinal assessments for RT, refraction measurements and VA were conducted postoperatively at 1 week, 1 month, 3 months, 6 months and then every 6 months thereafter (Figure 1c). The final VA was included in the analysis. Each type of examination was conducted by a single experienced examiner who was blind to the results of previous assessments to minimize potential bias. None of the assessments were mandatory when the infants were uncooperative or showed poor compliance, and these missing data were excluded. All the available data were included into analysis to ensure the fair representation of our study population (Table S1).

### Retinal development during initial stage blindness

To determine whether the retina shows responsiveness to external environment influences following initial stage blindness, we first used spectral domain optical coherence tomography (SD-OCT) to measure the RT of our patients at surgery. Ten healthy retinas as controls group were measured using the same procedures. A total of 56 retinas in cataract group (mean age, 3.5 months; range from 2 to 8 months) and 10 healthy retinas in control group (mean age, 4 months; range from 3 to 8 months) were ultimately included in this analysis.

The results revealed that the full-layer RT was thicker in the patients than in the control group during the initial stage blindness (5 retinal areas, *P*=0.012), and the differences were significant in the central fovea of the macula (Full layer in area 3: cataract vs. normal: 201.86±22.12 vs. 176.30±13.39, *P*=0.002). Moreover, the results also showed a thickening tendency of the inner layer of the retina of cataract group (5 retinal areas, P=0.036) and the differences were significant in the central fovea of the macula (Inner layer in area 3: cataract vs. normal: 65.43±12.45 vs. 55.70±5.59, *P*=0.026) (Figure 2a). The inner retina layer, containing the nerve fiber layer and ganglion cell layer, may be responsible for functional responses during the initial stage blindness.

**Figure 2.**
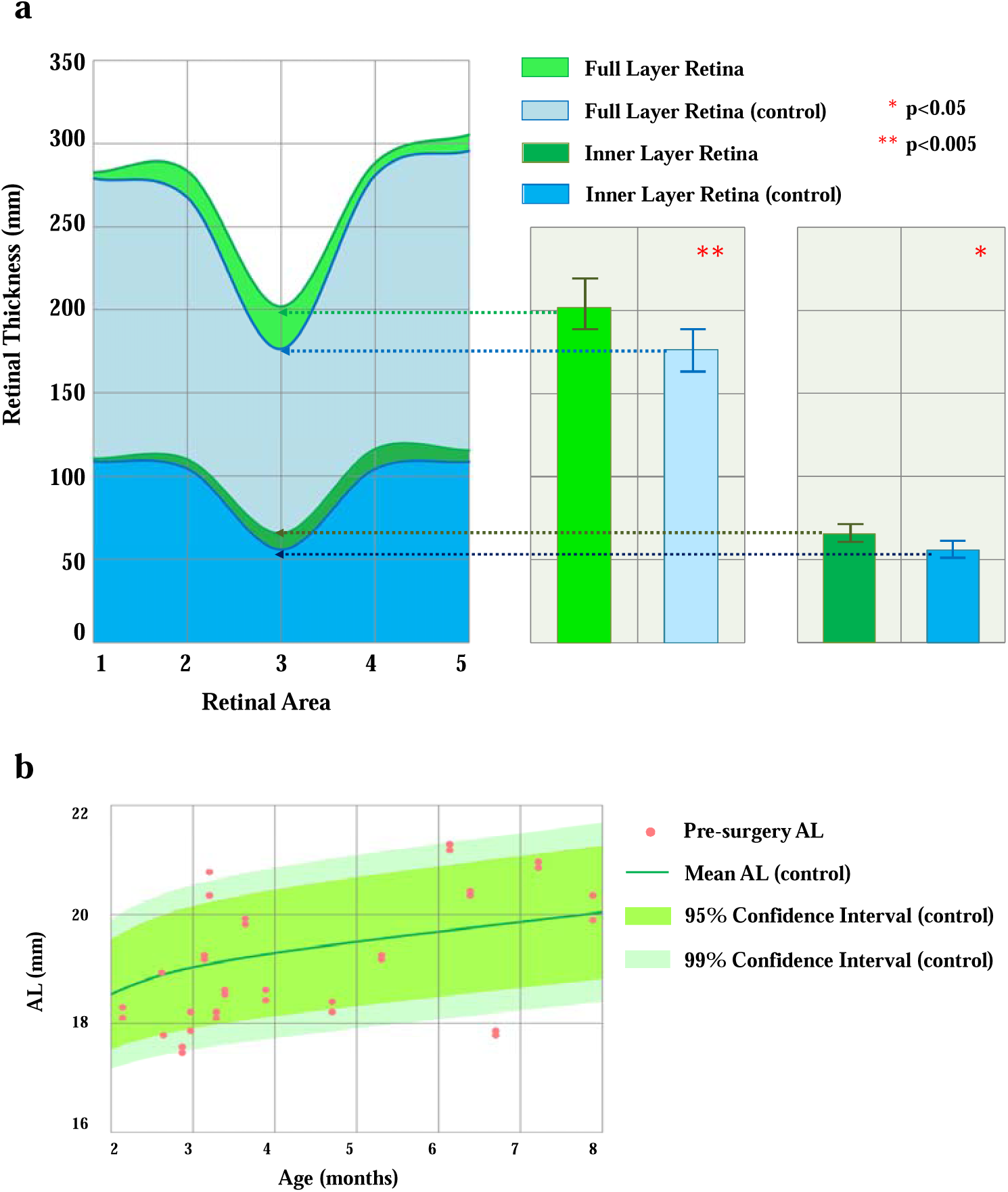
RT during surgery and pre-surgery axis length of cataract eyes compared with those of normal controls. **a,** The RT of cataract eyes (full layer and inner layer in retinal area 3, central fovea of the macula) were compared to those of normal controls and were found to be thicker in our patients. **b,** The control reference values were used to generate the normal AL range. The pre-surgery AL of our population was distributed mainly in the normal curve range; therefore, the AL development in our sample was considered similar to the normal level. (Notes: Bar graphs represent standard deviation; RT=retinal thickness; retina area 1=temporal peripheral area; 2=temporal para-central area; 3=central fovea of the macula; 4=nasal para-central area; 5=nasal peripheral area; AL=axial length).

### Retinal dynamic development following initial stage blindness

Then, we further investigate the dynamic changes in RT after surgery following the initial stage blindness. Four of the representative patients available with continuous follow-up records are presented (Figure 3a). As shown in Figure 3b, RT of our patients all experienced a slight decrease during the first week after surgery and a tendency to continuously increase during the following year. The dynamic RT changes during the first postoperative week might be mainly caused by the onset of vision while the continuously developmental tendency later demonstrates the retinal malleability.

**Figure 3.**
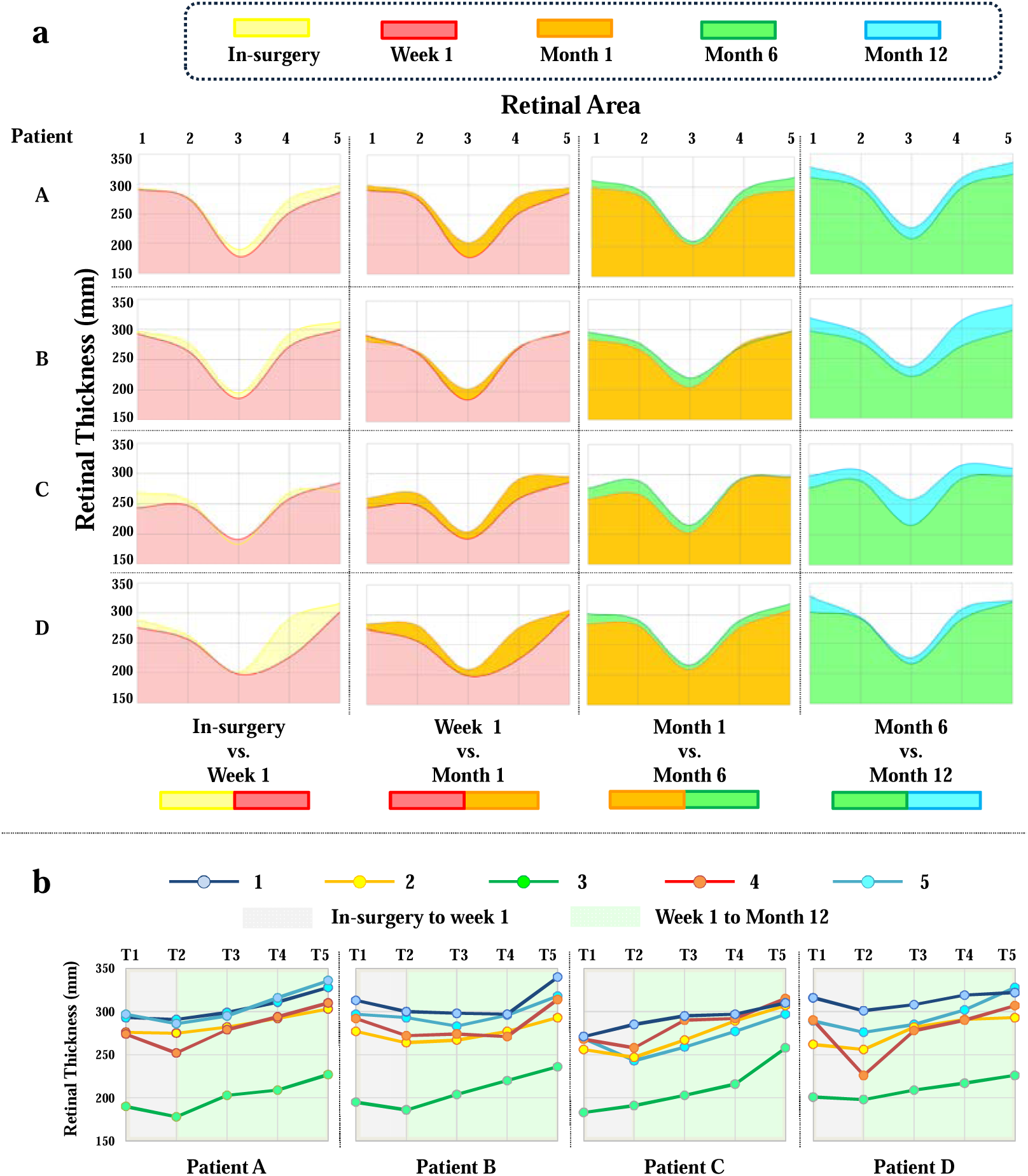
Dynamic developmental pattern of the retina following initial stage blindness. **a,** Continuous retinal changes of four representative patients with complete RT follow-up records are presented. During the first week after surgery, the RT underwent a slight thinning. From postoperative week 1 to month 12, the RT exhibited dynamic development. **b,** Developmental trend of RT is presented for each retinal area. (Notes: RT=retinal thickness; retina area 1=temporal peripheral area; 2=temporal para-central area; 3=central fovea of the macula; 4=nasal para-central area; 5=nasal peripheral area; T1=in-surgery; T2=week 1; T3=month 1; T4=month 6; T5=month 12).

We then evaluated the long-term retinal development by comparing the RT acquired during surgery with the RT at last follow up among all patients with available records. As shown in Figure 4a, all subjects showed a substantial RT increase at last follow up (mean age, 33 months; range from 20 to 49 months). Meanwhile, the RT in a group of 14 healthy retinas (mean age, 36 months; range from 21 to 47 months) was measured for comparison (Figure 4b). The RT values of the patients showed no significant differences with those of the control group in 5 retinal areas (*P*=0.77) and in each retinal area (5 regions from temporal to nasal respectively, cataract vs. control: 274.25±21.62 vs. 266.64±18.05, *P*=0.41; 280.13±20.52 vs. 291.29±16.62, *P*=0.20; 224.13±14.62 vs. 231.61±30.26, *P*=0.53; 299.75±14.46 vs. 302±23.28, *P*=0.81; 302.88±14.71 vs. 297.93±27.77, *P*=0.66). These results indicated that, although our patients exhibited individual differences in the growth of the retina throughout the longitudinal assessment, the retinal development of our patients ultimately reached a normal level.

**Figure 4.**
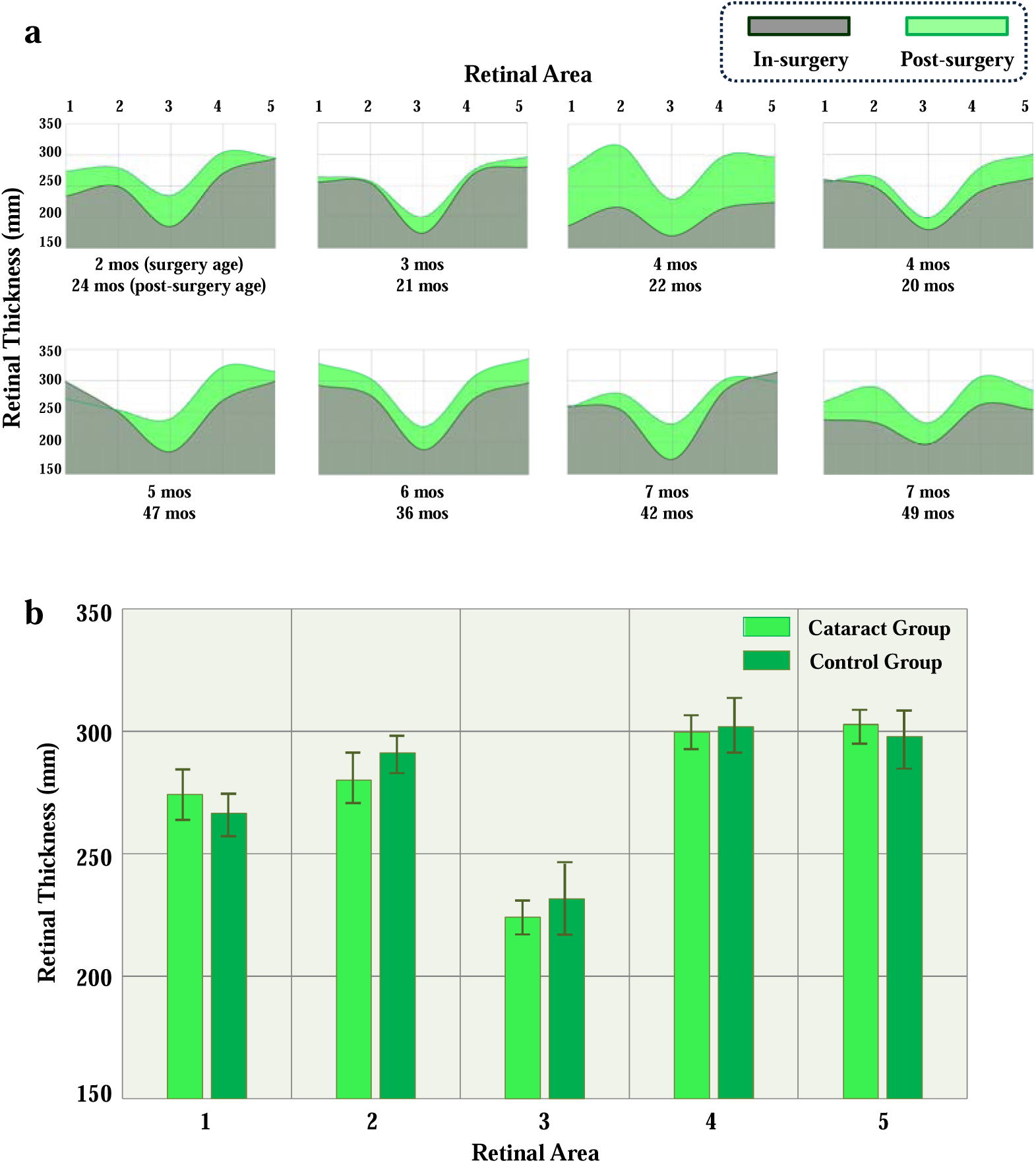
Endpoint development of the retina in patients compared with that in normal controls. **a,** The RT at the last follow up (green) was compared to the baseline RT during surgery (grey). Increases of RT were observed in all representative patients. **b,** No significant differences of RT in all five retinal area were observed between the patients and control groups. (Notes: Bar graphs represent standard deviation; mos=months; retina area 1=temporal peripheral area; 2=temporal para-central area; 3=central fovea of the macula; 4=nasal para-central area; 5=nasal peripheral area).

### AL development

We used all the available pre-surgery AL data from 30 eyes to investigate holistic eye emmetropization during initial stage blindness. To determine the normal rate of AL development, the referenced curve-fitting value of the age-matched normal distribution range was used for comparison (1 month: 17.00±0.40, 95%CI: 16.916-18.484, 99%CI: 16.670-18.730; 3 months: 19.03±0.58, 95%CI: 17.893-20.167, 99%CI: 17.536-20.524; 9 months: 20.23±0.64, 95%CI: 18.976-21.484, 99%CI: 18.581-21.879) (Pennie et al., 2001; Mutti et al., 2005). As shown in Figure 2b, the pre-surgery AL of our population was distributed mainly in the normal curve range. Therefore, the AL development before the onset of vision in our samples was considered to be similar to the normal level.

### Refractive dynamic development following initial stage blindness

Refractive status was evaluated following the onset of vision. All of the refractive changes of 64 eyes are presented in Figure 5a. Normal emmetropization of refractive media is generally considered to be 3 to 6 diopters during the first four years after birth. Refractive changes less than 3 diopters are considered undergrowth, whereas refractive changes of more than 6 diopters are considered overgrowth (Liu et al., 2016). The results showed that the majority of our patients (54 eyes, 84.4%) exhibited normal refractive development following initial stage blindness.

**Figure 5.**
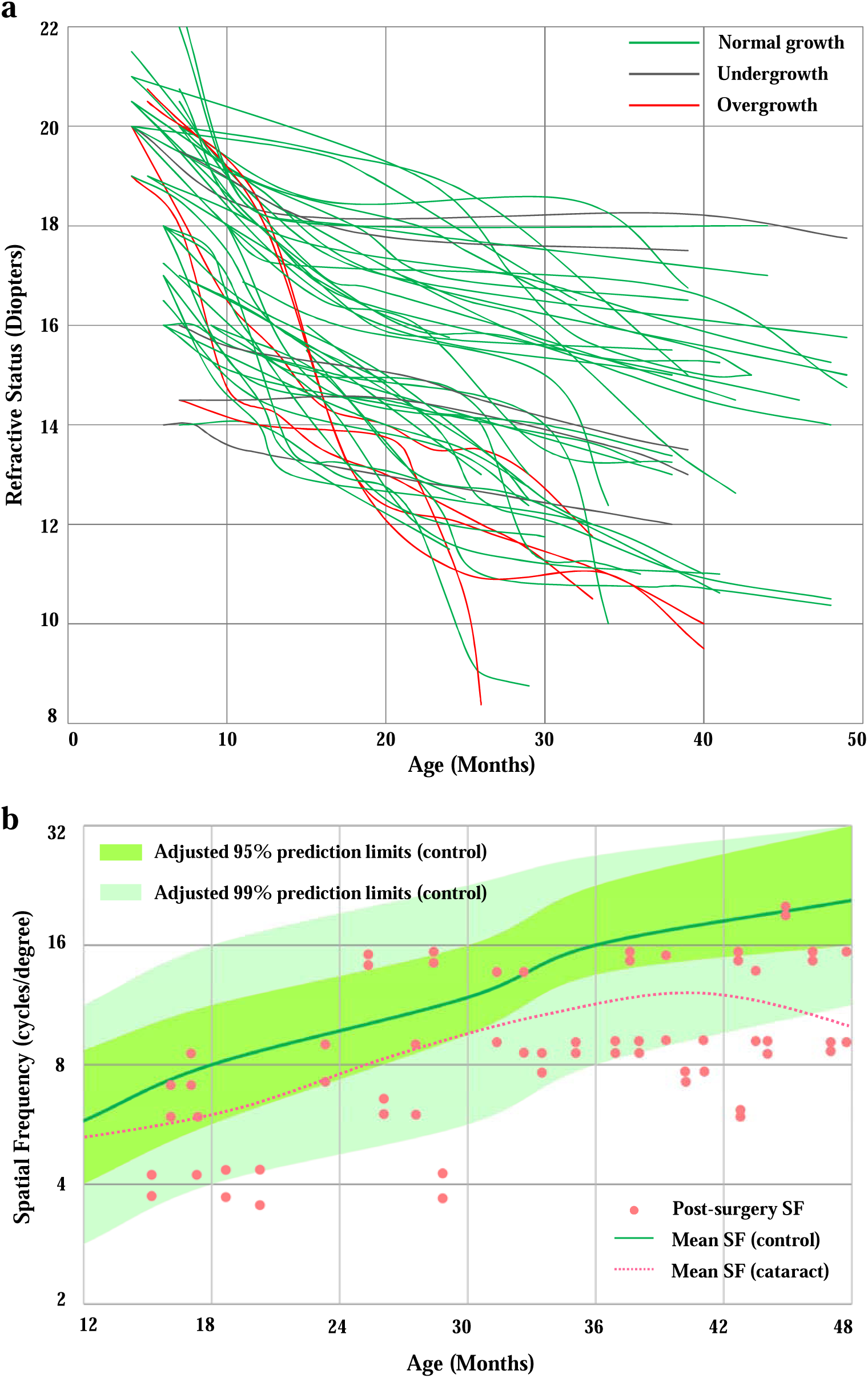
Dynamic developmental pattern of the refractive status and visual grating acuity following initial stage blindness. **a,** All refractive changes are presented. Refractive changes less than 3 diopters are considered undergrowth, refractive changes between 3 diopters to 6 diopters are considered normal growth, and refractive changes more than 6 diopters are considered overgrowth. The majority of our patients (54 eyes, 84.4%) exhibited normal refractive development following initial stage blindness. **b,** A normal distribution of the monocular grating acuity was referenced to evaluate our participants’ visual acuity. Our subjects showed improvements observed in visual acuity and the mean acuity of our patients is below the normal mean and falls outside of 95% prediction limits from around 2 years of age.

### Visual grating acuity assessment

To ensure the overall visual functional development of our population, Teller VA cards were used to assess the visual grating acuity of 60 eyes after surgery. The final visual acuity was used for the analysis. A normal distribution of monocular grating acuity and a referenced prediction limit were used (Mayer et al., 1995). Our subjects showed improvements observed in visual acuity (Figure 5b) and the mean acuity of our patients is below normal mean value and begins to fall outside the normal range around 2 years of age (Figure 5c).

### Genetic integrity of visual system development

Visual system development and maturation should be considered in the context of interactions between the environment and heredity. All of the included patients had no family heredity. Furthermore, no similar disease history (amblyopia or visual dysplasia) was observed in their immediate family members. Sampling investigation using whole exome sequencing was conducted for 7 children and their parents to confirm whether these children had the genomic deficiencies for visual system development. We sequenced the coding regions and all exon-intron boundaries for the 1679 known genes associated with human visual development. However, we found no direct relationships between filtered mutations and visual impairment according to the standard guidelines for the interpretation of sequence variants (Richards et al., 2015).

## DISCUSSION

Visual experience is thought to mediate and drive visual system development. Infants are born with rudimentary visual capabilities and require sufficient visual experience early in life to reach optimal levels of visual functioning as adults (Lewis and Maurer, 2009). However, each year, millions of infants worldwide suffer from visual deprivation. These populations face the risk of irreversible amblyopia and numerous vision impairments (d’Almeida et al., 2013; Mansouri et al., 2013; Lin et al., 2016). Thus, it is important to investigate the mechanisms of environmental regulation of visual system development and its experience-dependent plasticity, which may provide further evidence to develop a comprehensive method for assessing the visual recovery potential in blind children.

Various components of the eyeball have been shown to undergo periods of experience-dependent development, with evidences from both human and non-human animal experiments indicating that prolonged deprivation of form vision leads to increased AL and myopia (Fledelius et al., 2014; Lin et al., 2016). Moreover, initial stage blindness also influences the functional and morphological maturation of the retina, including its synaptic density and bipolar cell structure (Tian and Copenhagen, 2001). However, little is known about how these diverse parts of the visual system are interrelated and interact with each other.

In summary, our findings demonstrate that retina is malleable and associated with external environment influences. Laties AM and colleagues once posited that the retina may participate in the postnatal regulation of eye growth to minimize refractive error (Stone et al., 1990; Laties AM, 1991). Recent studies have shown that the axial overgrowth and myopia caused by visual form deprivation can be manipulated by altering peripheral retinal defocus (Benavente-Pérez et al., 2014). In addition, early retinal changes are reflected in retinotopically specific plasticity, which can be assessed by visual cortical thickness (d’Almeida et al., 2013; Mateus et al., 2016). Both neurochemical and immunocytochemical experiments in chickens and monkeys suggest that definable retinal neurons participate in the regulatory pathway controlling eye growth (Laties AM, 1991). All these lines of evidence suggest that the retina may act as an intact connection to the anterior segment optic system as well as the visual cortex during early visual development (sketch map shown in Figure 6a). The “bridge” role of the retina may be functionally consistent with that of dopamine receptors, which are thought to regulate synapse formation, synaptic transmission, and light adaptation in the experience-dependent development of the retina (He et al., 2013; Tian et al., 2015).

**Figure 6.**
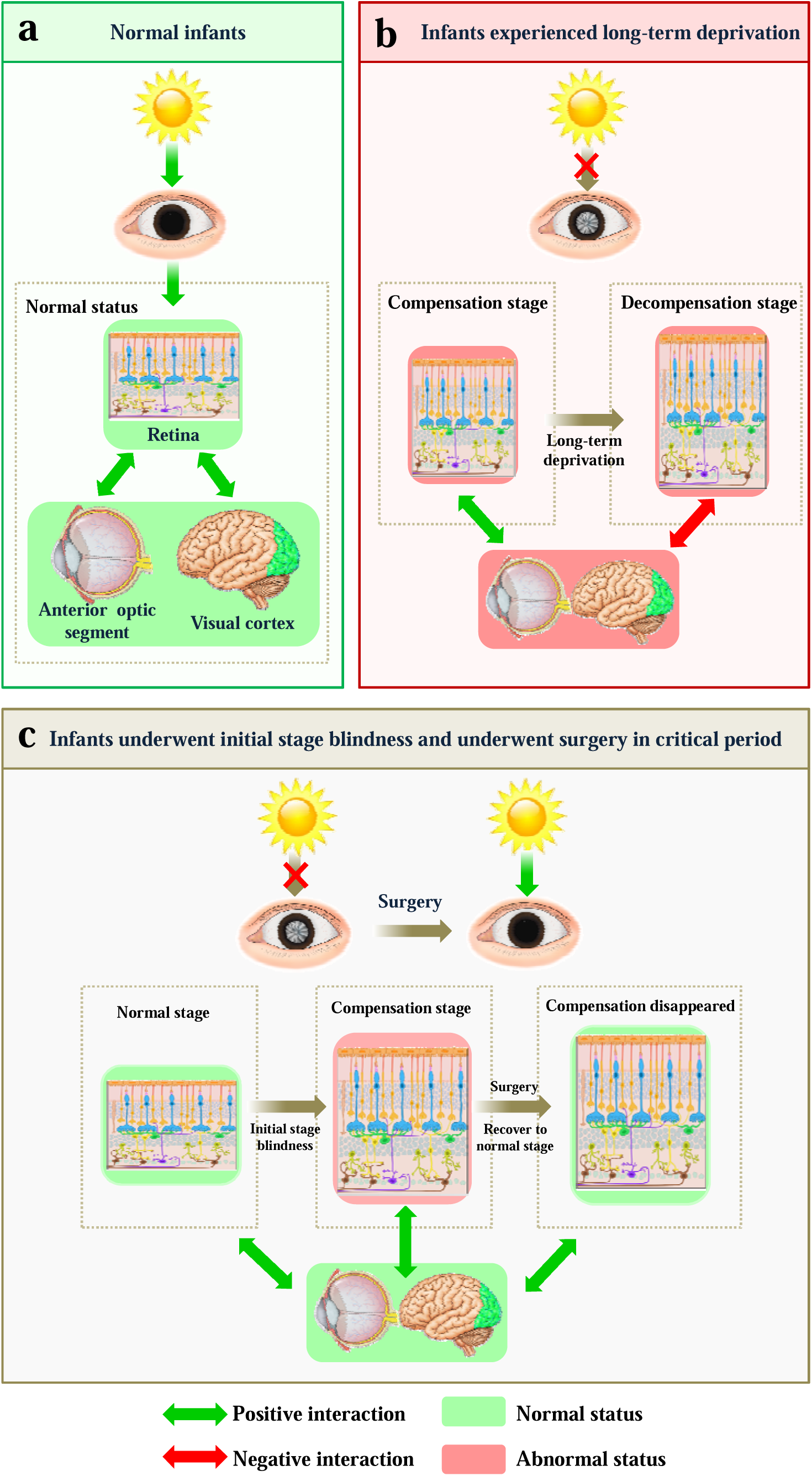
The retina plays a crucial role in visual system development and the dynamic changes of RT might account for sensitive period plasticity in humans. **a,** The retina acts as a bridge connecting the external environment and visual system components from the anterior optic segment to the visual cortex. **b,** The retina will extend to the decompensation stage when experiencing long-term deprivation, accompanied with the anterior optic segment and visual cortex undergoing fluctuating changes, thus lead to irreversible vision impairment. **c,** The retina accompanied with other visual system parts, has a latent thickening tendency during initial stage blindness to functionally compensate for the insufficient visual stimulations. As soon as the external signals reach the retina and visual system successfully (after surgery), the compensation disappeared and returned to the normal developmental tendency.

It is well known that during sensitive periods, the visual system is vulnerable to the harmful effects of deprivation but still has the potential to recover. This recovery potential, called plasticity, is a crucial factor in establishing mature circuitry (Hooks and Chen, 2007). We found that the retina has a latent thickening tendency during initial stage blindness, which might be presumed to reflect an attempt to functionally compensate for the insufficient visual stimulation and to prepare for the potential following signal penetration (sketch map shown in Figure 6c). Previous studies indicated that during initial stage blindness the increasing expression of amacrine cells is triggered, with nerve growth factors and brain-derived neurotropic factors also involved to induce retinal light adaptation and contrast enhancement (Kim and von, 2016). All these functional responses might be involved in our dynamic procedure. After surgery, retinal compensation was found to be disappeared, and the development of retina was gradually recovering. This commutation activity of the retina might partly explain its recovery potential during sensitive periods. Retina may presumably extend to the decompensation stage during long-term visual deprivation accompanied with abnormal development of anterior segment optic system and visual cortex, thereby leading to irreversible visual impairments (sketch map shown in Figure 6b).

Our study has three implications. First, the visual system development should be considered as a whole, with the retina acting as a bridge that connects the external environment with each visual system component, from the anterior segment optic system to the visual cortex. Second, the intrinsic reason that accounts for visual plasticity might be a compensation process, as the dynamic changes of RT in our study reflect functional adaptation in response to the initial stage blindness. Third, we tentatively propose that RT might be used as a direct and sensitive indicator of abnormal visual stimulation as well as plasticity, which may provide an opportunity to develop a novel method for assessing visual recovery potential in blind children.

The results of our study should be cautiously interpreted within the context of two main limitations. First, our study primarily measures the effects of initial stage blindness on the retina but not the brain. Vision is a collaborative function of the retina and the brain. Therefore, dramatic changes of visual cortex correlated with the retina might be detected if the brain was investigated as well, which might account for the reason why part of our patients have lags in visual function development. Second, we used two control groups for the comparison of RT at surgery and at the last follow up. Although an age and number matched parallel group of control is a better choice for comparison, measuring the retinal thickness using SD-OCT is not necessary for a healthy child and therefore it is impractical to set a parallel control group for such a long-term follow-up study.

Previous studies reported that the fellow eye of unilateral congenital cataract patients, which likely has a normal retina, shows deficits in various aspects of vision (Lewis et al. 1992). Therefore, potential factors including biased interocular competition might influence the plasticity as well, which remains to be investigated in the future. Moreover, additional long-term and complete records are needed to investigate the effect of age, which could explain why the VA of our patients begins to fall outside of normal range around 2 years of age, as reported in a previous study (Lewis et al. 1995). Meanwhile, future researches on visual cortex examination and additional measures for peripheral retinal thickness will provide further understanding of visual system development.

## MATERIALS AND METHODS

### Study population

In total, thirty-nine individuals registered with the Childhood Cataract Program of the Chinese Ministry of Health (CCPMOH) (Lin et al., 2015) were recruited between January 2010 and March 2011 from Zhongshan Ophthalmic Center (ZOC), one of the largest eye hospitals in China (Dolgin, 2015). All participants were born with dense and total bilateral cataracts, diagnosed (mean age, 2.9 months; range from 1 to 7.5 months) and underwent surgery for bilateral cataract removal (mean age, 3.5 months; range from 2 to 8 months) at an early age. The first prescription of glasses was assigned to the participants at 1 week after surgery. All the prescription changes in glasses were decided by experienced optometrists. Our participants completed their follow-ups at mean age of 37.8 months, ranged from 20 to 49 months. Apart from their history of cataracts, all individuals were healthy (e.g., no metabolic diseases, mental retardation or central nervous diseases) and had no history of inherited diseases.

### Ethical approval

The research protocol was approved by the Institutional Review Board/Ethics Committee of Sun Yat-sen University (Guangzhou, China). Informed written consent was obtained from at least one family member of each participating child, and the tenets of the Declaration of Helsinki were followed throughout the study. To allow confidential evaluation using a slit-lamp, a SD-OCT imaging system, an A scan, a retinoscopy and the Teller VA card during our study, this trial was registered with the Clinical Research Internal Management System of ZOC.

### IVue OCT for RT measurements

We used an SD-OCT system (iVue SD-OCT; Optovue, Inc., Fremont, CA, USA) to evaluate RT. The protocol of iVue OCT consists of 12 radial scans of 3.4 mm in length (452 A scans each) and 6 concentric ring scans ranging from 2.5 to 4.0 mm in diameter (587 to 775 A scans each), all centered on the optic disc. All of the images were reprocessed with a three-dimensional/video baseline. The parameters measured by the software included the optic disc, optic cup, neuroretinal rim, nerve head volume, cup volume, rim volume, cup-disc area ratio, horizontal cup-disc ratio, and vertical cup-disc ratio. The protocol also generates a polar thickness map, measured along a circle of 3.45 mm in diameter and centered on the optic disc. The procedure provides the average in the temporal, superior, nasal, inferior quadrants and the overall average along the entire measurement circle. The peripheral, para-central and central RTs from the temporal to nasal area were used here in the final analysis.

### A scan for AL measurements

Before surgery, a contact A scan (B-SCAN-Vplus /BIOVISION, Quantel Medical, France) was used for AL measurements. The A scan unit was equipped with a 10 MHz transducer probe, and the velocities were set as follows: 1,641 m/s for the cornea and lens and 1,532 m/s for the aqueous and vitreous humor. Applanation ultrasound was performed after the instillation of one drop of topical anesthetic (0.5% Alcaine, Alcon, USA) to the lower conjunctiva. Each eye was measured 10 times, and the mean measurements were used for the final analysis.

### Refraction and VA measurement

All refractions were conducted using objective retinoscopy and cycloplegia. The spherical equivalent power was included in the analysis. All of the monocular best-corrected visual grating acuity was measured with glasses using a complete set of Teller VA Cards (Stereo Optical Company, Inc., IL, USA) (Mayer et al., 1995). The set consisted of 15 cards with gratings ranging in spatial frequency from 0.32 cycles/cm to 38 cycles/cm in half-octave steps as well as a low vision card and a blank gray card. Luminance was kept above 10 candelas/m^2^ by utilizing overhead diffuse fluorescent lighting and a spotlight directed towards the ceiling; in addition, the contrast of the cards is approximately 60-70%. Infants were assessed according to a standard procedure in the operation manual (Cavallini et al., 2002; Ciocler and Dantas, 2013). The order of testing eyes (right/left) was randomized across children.

### Exome-capture sequencing and variant calling

Genomic DNA was extracted from blood using a QIAGEN DNeasy Blood and Tissue Kit (QIAGEN, USA) according to the manufacturer’s protocol. Isolated genomic DNA from blood was captured by Roche’s Nimblegen SeqCap EZ Human Exome v2.0 library using in-solution hybridization and PCR to enrich the exomes before sequencing. Illumina HiSeq X10 was used to perform next-generation sequencing to evaluate differences in mutations. The sequencing reads of each sample were aligned to the human reference genome hg19 assembly using Burrows-Wheeler Aligner (Li and Durbin, 2009), SAMtools and Picard tools. The 1679 known genes associated with human visual development were collected for genetic analysis (Table S2). The snps and indels were detected by HaplotypeCaller according to the instructions. ANNOVAR were used to annotate all the variants. Variants with a frequency more than 1% in dbSNP, 1000 genome, ESP6500 or the in-house database were excluded. PolyPhen-2, SIFT and MutationTaster were used to predict the effect protein function of amino acid substitution. In addition to de novo mutations, compound heterozygous mutations and homozygous mutations were considered based on the recessive model. However, we found no direct relationships between filtered mutations and visual impairment according to the standards and guidelines for the interpretation of sequence variants. (Richards et al., 2015)

### Statistical analysis

Mixed ANOVA were used to compare RT differences (5 retinal areas) between cataract and control groups. An independent-sample t-test was used to compare RT differences between cataract and control groups in each retinal area. The Bonferroni method was used to correct alpha for multiple t-test (α’= α/m, α = 0.05 and m is the number of hypotheses). All statistical tests were two-tailed, and a *P*-value below 0.05 or corrected alpha was considered statistically significant. All statistical analyses were performed using SPSS software, v. 18 (SPSS, Inc., Chicago, IL, USA).

## ACKNOWLEDGMENTS

The authors would like to thank volunteer statistical expert, Zicong Chen, for discussions and critical reading of the manuscript.

## AUTHOR CONTRIBUTIONS

H.T.L., E.P.L. and X.Y.Z. designed the study; E.P.L., X.Y.Z., Z.Z.L., X.H.W., X.H.T., D.R.L., Q.Z.C., J.J.C., Z.L.L., X.Y.L., J.L., D.N.W., J.H.W., W.T.L., L.X.L., W.R.C. and Y.Z.L. performed the research. E.P.L. and X.Y.Z. analyzed the data. H.T.L., E.P.L. and X.Y.Z. co-wrote the manuscript, and all authors discussed the results and commented on the paper.

## Supplementary Information

**Table S1. Overview of the clinical records for all included patients.**

**Table S2. Known genes associated with human visual development.**

## REFERENCE

Azuma, N., Yamaguchi, Y., Handa, H., Hayakawa, M., Kanai, A. and Yamada, M. (1999). Missense mutation in the alternative splice region of the PAX6 gene in eye anomalies. Am. J. Hum. Genet. 65, 656-663.

Barbieri, A.M., Lupo, G., Bulfone, A., Andreazzoli, M., Mariani, M., Fougerousse, F., Consalez, G.G., Borsani, G, Beckmann, J.S., Barsacchi, G., et al. (1999). A homeobox gene, vax2, controls the patterning of the eye dorsoventral axis. Proc. Natl. Acad. Sci. U.S.A. 96, 10729-10734.

Benavente-Pérez, A., Nour, A. and Troilo, D. (2014). Axial eye growth and refractive error development can be modified by exposing the peripheral retina to relative myopic or hyperopic defocus. Invest. Ophthalmol. Vis. Sci. 55, 6765-6773.

Cavallini, A., Fazzi, E., Viviani, V., Astori, M.G., Zaverio, S., Bianchi, P.E. and Lanzi, G. (2002). Visual acuity in the first two years of life in healthy term newborns: an experience with the teller acuity cards. Funct. Neurol. 17, 87-92.

Ciocler, F.P. and Dantas, P.E.. (2013). Assessment of visual acuity in patients with dementia using teller acuity cards. Strabismus 21, 93-97.

d’Almeida, O.C., Mateus, C., Reis, A., Grazina, M.M. and Castelo-Branco, M. (2013). Long term cortical plasticity in visual retinotopic areas in humans with silent retinal ganglion cell loss. Neuroimage 81, 222-230.

de Heering A, Dormal G, Pelland M, Lewis T, Maurer D, Collignon O. (2016) A Brief Period of Postnatal Visual Deprivation Alters the Balance between Auditory and Visual Attention. Curr Biol. Nov 21;26(22):3101-3105.

Dolgin, E. (2015). The myopia boom. Nature 519, 276-278.

Fledelius, H.C., Goldschmidt, E., Haargaard, B. and Jensen, H. (2014). Human parallels to experimental myopia? A literature review on visual deprivation. Acta Ophthalmol 92, 724-729.

Grady, C.L., Mondloch, C.J., Lewis, T.L., and Maurer, D. (2014). Early visual deprivation from congenital cataracts disrupts activity and functional connectivity in the face network. Neuropsychologia 57, 122-139.

Guerreiro MJ, Putzar L, Röder B. (2016). Persisting Cross-Modal Changes in Sight-Recovery Individuals Modulate Visual Perception. Curr Biol. Nov 21;26(22):3096-3100.

Hallonet, M., Hollemann, T., Pieler, T. and Gruss, P. (1999). Vax1, a novel homeobox-containing gene, directs development of the basal forebrain and visual system. Genes Dev. 13, 3106-3114.

He, Q., Xu, H.P., Wang, P. and Tian, N. (2013). Dopamine D1 receptors regulate the light dependent development of retinal synaptic responses. PLoS ONE 8, e79625.

Hooks, B.M. and Chen, C. (2007). Critical periods in the visual system: changing views for a model of experience-dependent plasticity. Neuron 56, 312-326.

Jeon ST, Maurer D and Lewis TL. (2012). The effect of video game training on the vision of adults with bilateral deprivation amblyopia. Seeing Preceiving 25, 493-520.

Kim, M.H. and von, G.H. (2016). Postsynaptic Plasticity Triggered by Ca^2+^’-Permeable AMPA Receptor Activation in Retinal Amacrine Cells. Neuron 89, 507-520.

Laties AM, S.R.A. (1991). Some visual and neurochemical correlates of refractive development. Vis Neurosci. Jul-Aug;7(1-2), 125-128.

Lewis, TL, Maurer D, et al. (1992). Vision in the “good” eye of children treated for unilateral congenital cataract. Ophthalmology. Jul; 99(7):1013-7.

Lewis, TL, Maurer D, et al. (1995). Dvelopment of grating acuity in children treated for unilateral or bilateral congenital cataract. Invest Ophthalmol Vis Sci. Sep; 36(10):2080-95.

Lewis, T.L. and Maurer, D. (2005). Multiple sensitive periods in human visual development: evidence from visually deprived children. Dev Psychobiol 46, 163-183.

Lewis, T.L. and Maurer, D. (2009). Effects of early pattern deprivation on visual development. Optom Vis Sci 86, 640-646.

Li, H. and Durbin, R. (2009). Fast and accurate short read alignment with Burrows-Wheeler transform. Bioinformatics 25, 1754-1760.

Lin, H., Lin, D., Chen, J., Luo, L., Lin, Z., Wu, X., Long, E., Zhang, L., Chen, H., Chen,W., et al. (2016). Distribution of Axial Length before Cataract Surgery in Chinese Pediatric Patients. Sci Rep 6, 23862.

Lin, H., Long, E., Chen, W. and Liu, Y. (2015). Documenting rare disease data in China. Science 349, 1064.

Lin, H., Ouyang, H., Zhu, J., Huang, S., Liu, Z., Chen, S., Cao, G., Li, G., Signer, R.A., Xu, Y., et al. (2016). Lens regeneration using endogenous stem cells with gain of visual function. Nature 531, 323-328.

Liu, Y., Yu, C., Liang, M., Li, J., Tian, L., Zhou, Y., Qin, W., Li, K. and Jiang, T. (2007). Whole brain functional connectivity in the early blind. Brain 130, 2085-2096.

Liu, Z.. Long, E., Chen, J., Lin, Z., Lin, D., Wu, X., Cao, Q., Li, X., Wang, D., Luo, L., et al. (2016). Developmental profile of ocular refraction in patients with congenital cataract: a prospective cohort study. The Lancet. October 388:S54.

Lundwall, R.A., Dannemiller, J.L. and Goldsmith, H.H. (2015). Genetic associations with reflexive visual attention in infancy and childhood. Dev Sci

Mansouri, B., Stacy, R.C., Kruger, J. and Cestari, D.M. (2013). Deprivation amblyopia and congenital hereditary cataract. Semin Ophthalmol 28, 321-326.

Mateus, C., d’Almeida, O.C., Reis, A., Silva, E. and Castelo-Branco, M. (2016). Genetically induced impairment of retinal ganglion cells at the axonal level is linked to extrastriate cortical plasticity. Brain Struct Funct 221, 1767-1780.

Maurer, D., Lewis, T.L., Brent, H.P. and Levin, A.V. (1999). Rapid improvement in the acuity of infants after visual input. Science 286, 108-110.

Mayer, D.L., Beiser, A.S., Warner, A.F., Pratt, E.M., Raye, K.N. and Lang, J.M. (1995). Monocular acuity norms for the Teller Acuity Cards between ages one month and four years. Invest. Ophthalmol. Vis. Sci. 36, 671-685.

Minoshima, S., Mitsuyama, S., Ohno, S., Kawamura, T. and Shimizu, N. (1999). Keio Mutation Database for eye disease genes (KMeyeDB). Nucleic Acids Res. 27, 358-361.

Morishita, H. and Hensch, T.K. (2008). Critical period revisited: impact on vision. Curr. Opin. Neurobiol. 18, 101-107.

Mutti, D.O., Mitchell, G.L., Jones, L.A., Friedman, N.E., Frane, S.L., Lin, W.K., Moeschberger, M.L. and Zadnik, K. (2005). Axial growth and changes in lenticular and corneal power during emmetropization in infants. Invest. Ophthalmol. Vis. Sci. 46, 3074-3080.

Mutti, D.O., Mitchell, G.L., Jones, L.A., Friedman, N.E., Frane, S.L., Lin, W.K., Moeschberger, M.L. and Zadnik, K. (2009). Accommodation, acuity, and their relationship to emmetropization in infants. Optom Vis Sci 86, 666-676.

Pennie, F.C., Wood, I.C., Olsen, C., White, S. and Charman, W.N. (2001). A longitudinal study of the biometric and refractive changes in full-term infants during the first year of life. Vision Res. 41, 2799-2810.

Qin, W., Liu, Y., Jiang, T. and Yu, C. (2013). The development of visual areas depends differently on visual experience. PLoS ONE 8, e53784.

Richards, S., Aziz, N., Bale, S., Bick, D., Das, S., Foster, G.J., Grody, W.W., Hegde, M., Lyon, E., Spector, E., et.al. (2015). Standards and Guidelines for the Interpretation of Sequence Variants: A Joint Consensus Recommendation of the American College of Medical Genetics and Genomics and the Association for Molecular Pathology. Genetics in medicine: official journal of the American College of Medical Genetics. 17, 405-423.

Rolle, T., Manerba, L., Lanzafame, P. and Grignolo, F.M. (2016). Diagnostic Power of Macular Retinal Thickness Analysis and Structure-Function Relationship in Glaucoma Diagnosis Using SPECTRALIS OCT. Curr. Eye Res. 41, 667-675.

Schaeffel, F. and Wildsoet, C. (2013). Can the retina alone detect the sign of defocus. Ophthalmic Physiol Opt 33, 362-367.

Skoczenski, A.M. and Norcia, A.M. (1999). Development of VEP Vernier acuity and grating acuity in human infants. Invest. Ophthalmol. Vis. Sci. 40, 2411-2417.

Smith, E.L., Hung, L.F. and Arumugam, B. (2014). Visual regulation of refractive development: insights from animal studies. Eye (Lond) 28, 180-188.

Stone, R.A., Lin, T., Iuvone, P.M. and Laties, A.M. (1990). Postnatal control of ocular growth: dopaminergic mechanisms. Ciba Foundation symposium 155, 45-57; discussion 57-62.

Tian, N. and Copenhagen, D.R. (2001). Visual deprivation alters development of synaptic function in inner retina after eye opening. Neuron 32, 439-449.

Tian, N., Xu, H.P. and Wang, P. (2015). Dopamine D2 receptors preferentially regulate the development of light responses of the inner retina. Eur. J. Neurosci. 41, 17-30.

Wallman, J. and Winawer, J. (2004). Homeostasis of eye growth and the question of myopia. Neuron 43, 447-468.

Wang, X., Peelen, M.V., Han, Z., He, C., Caramazza, A. and Bi, Y. (2015). How Visual Is the Visual Cortex? Comparing Connectional and Functional Fingerprints between Congenitally Blind and Sighted Individuals. J. Neurosci. 35, 12545-12559.

